# Temporal Lobe Epilepsy is dominated by Region Specific Interictal Cortical Inhibition

**DOI:** 10.1101/2024.11.30.626179

**Authors:** Ricardo Kienitz, Adam Strzelczyk, Andrea Spyrantis, Jürgen Konczalla, Felix Rosenow, Michael Strüber

**Author notes:** Correspondence should be addressed to RK.

## Abstract

Epilepsy is typically characterized by excessive neuronal excitability, manifesting as seizures and interictal epileptiform discharges (IEDs) in the EEG. However, the dynamics of excitation and inhibition (E/I balance) remain poorly understood. Here, we leverage the aperiodic exponent of the EEG power spectrum—a marker indicative of synaptic inhibition—to investigate shifts in E/I balance during antiseizure medication (ASM) tapering in patients with mesial temporal lobe epilepsy (TLE). We analyzed EEG data from 28 TLE patients and 25 controls with non-epileptic episodes (NEE) undergoing presurgical video EEG monitoring. Unexpectedly, TLE patients showed a localized increase in the aperiodic exponent in the ipsilesional temporal lobe during ASM tapering, absent in controls. This inhibition increase correlated with seizure latency and predicted seizure occurrence. Intracranial recordings from 10 TLE patients revealed higher aperiodic exponents in the lateral temporal cortex compared to the hippocampus, suggesting stronger inhibition in the lateral cortex. Notably, hippocampal IEDs triggered transient inhibitory responses in the lateral cortex, accompanied by increased high-frequency activity and disrupted hippocampus-to-lateral connectivity. These findings suggest that TLE likely involves complex inhibitory mechanisms beyond the epileptic focus in the interictal period, with neocortical inhibition potentially containing epileptic activity, and offers a new tool to map epileptic brain regions.

## Introduction

Epilepsy is a chronic neurological disorder characterized by recurrent, unprovoked seizures, affecting approximately 1% of the global population (Ngugi et al. 2010). Mesial temporal lobe epilepsy (TLE) is among the most common focal and refractory forms of human epilepsy, with the hippocampus frequently serving as the primary epileptogenic zone.

Mechanistically, epilepsy is closely linked to abnormal neuronal excitability. Disruptions in the physiological balance between excitation and inhibition (E/I balance) are considered a key mechanism underlying epilepsy, where excessive excitation or insufficient inhibition can result in pathological hyperexcitability and seizure generation (Huberfeld et al. 2007; Staley 2015). For example, the hippocampus in mesial temporal lobe epilepsy often shows a loss of GABAergic interneurons or dysfunction in inhibitory synapses (Marx, Haas, and Häussler 2013; de Lanerolle et al. 1989), resulting in a downregulation of inhibitory activity.

Despite its critical importance, changes in the E/I balance are not readily accessible in electrophysiological macro-recordings. Standard clinical EEG markers, such as interictal epileptiform discharges (IEDs) and high-frequency oscillations (HFOs), have traditionally been interpreted as indicators of heightened excitability (Fisher et al. 1992). However, the dynamics and mechanisms of E/I balance in patients remain poorly understood, particularly during the interictal state, which constitutes the vast majority of a patient’s time. Unlike seizures, the interictal state further lacks behavioral markers, making it challenging to evaluate despite its profound clinical relevance.

Recent advances in EEG analysis techniques have provided new tools to quantify balance between excitation and inhibition (E/I balance) in vivo. One such method involves analyzing the aperiodic component of the EEG power spectrum, specifically the slope of the 1/f component, known as the aperiodic exponent (He 2014; Gao, Peterson, and Voytek 2017; Donoghue et al. 2020) A steeper slope is indicative of greater inhibition, offering a marker of synaptic E/I dynamics in both healthy and pathological states. Very recently, we identified changes in inhibitory tone following epileptogenic interventions in a TLE rat model. Here, the aperiodic exponent demonstrated a long-lasting increase following electrical stimulation mimicking status epilepticus and was predictive of later epilepsy development. Furthermore, a recent study further found larger aperiodic exponents in epilepsy patients compared to healthy controls (Duma et al. 2024).

In this study, we leveraged tapering of anti-seizure medication (ASM) as a clinical tool commonly performed during video EEG monitoring (VEM) to record seizures in epilepsy patients. As ASM tapering increases the likelihood of seizure occurrence it is commonly thought to increase excitability.

To test this, we first calculated the aperiodic exponent to quantify synaptic inhibition in standard EEG recordings from TLE epilepsy patients during VEM and ASM tapering. Patients with non-epileptic episodes only (NEE), who also underwent ASM tapering, served as controls. To gain further insight into the mechanisms of E/I dynamics, we next analyzed intracranial recordings from TLE patients, focusing on the hippocampus as the seizure onset zone (SOZ) in these patients and the lateral temporal cortex as a connected area outside of the SOZ which is also accessible to standard EEG. We investigated the regional differences in the E/I balance between these areas and explored the role of hippocampal IEDs in driving changes in inhibition within the lateral cortex.

Our findings challenge the traditional view that ASM tapering uniformly increases neuronal excitability across the brain. Instead, we observed a paradoxical increase in inhibition localized to the ipsilesional temporal lobe in epilepsy patients, while no such changes were observed in NEE controls. Importantly, we show that this phenomenon can be detected using non-invasive, surface EEG used for routine work-up of epilepsy patients, offering significant implications for diagnosing and mapping epileptic changes in these patients.

Intracranial recordings revealed that hippocampal IEDs drive transient increases in lateral cortical inhibition, accompanied by increased neuronal firing, as reflected by high-frequency band activity. Additionally, hippocampal IEDs disrupted the hippocampus’s dominant influence over the lateral cortex, shifting connectivity dynamics toward greater lateral-to-hippocampus influence. These results suggest that TLE involves complex, region-specific inhibitory mechanisms in the interictal period that may serve to limit the spread of epileptic activity.

Our findings provide new insights into the dynamics of excitation and inhibition in epilepsy, offering a deeper understanding of inhibitory mechanisms that could inform future studies on ASM effectiveness.

## Results

### Neuronal Complexity is reduced in epilepsy patients

We first recorded EEG to clinical standard in 28 patients with unilateral mesial temporal lobe epilepsy (TLE) undergoing presurgical Video-EEG monitoring (VEM) and tapering of anti-seizure medication (ASM) to increase the likelihood of seizures (**Figure 1A**). 25 patients with non-epileptic episodes only (NEE) who also underwent ASM tapering served as a control group.

**Figure 1.**
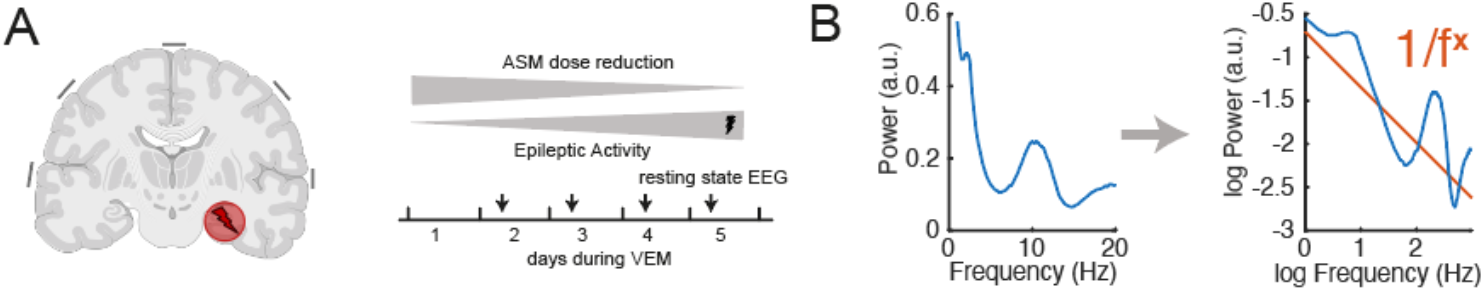
Experimental design and methodology for quantifying synaptic E/I balance. **(A)** Illustration of the study design. Patients with unilateral mesial temporal lobe epilepsy (TLE) were included. Right panel depicts the timeline of antiseizure medication (ASM) tapering during presurgical video-EEG monitoring (VEM). Patients with non-epileptic episodes only (NEE) served as controls. Patients received daily resting state EEG recordings. **(B)** Schematic of the EEG power spectrum analysis. The left panel shows the raw power spectrum, while the right panel illustrates the 1/f fit (orange) in log-log space. The slope of the 1/f component (aperiodic exponent) was used as a marker of synaptic excitation/inhibition (E/I) balance, computed from 3-second EEG segments across consecutive days of ASM tapering.

We hypothesized that ASM tapering during VEM would lead to increased neuronal excitability culminating in seizures. We therefore first aimed to assess changes of synaptic E/I balance over consecutive days. To this end, we computed the aperiodic exponent (Donoghue et al. 2020), which captures the slope of the 1/f component of the EEG’s powerspectrum (**Figure 1B**) and has recently been shown to quantify E/I balance in macrorecordings (Gao, Peterson, and Voytek 2017; see Ahmad et al. 2022, for a review). Daily EEG recordings were cut into 3s segments which served to compute the aperiodic exponent. Changes during ASM tapering were quantified as the mean percent change per day between the first resting state measurement and the day of minimum ASM dose.

When comparing fitted 1/fx functions between the initial and the minimum-ASM recording, we noted a prominent increase of the slope (see **Figure 2A** for one representative patient). This indicates an increase in the aperiodic exponent suggestive of an increase in inhibition. In contrast to our initial hypothesis, we found this increase of the aperiodic exponent to be significantly positive in TLE patients (p = 0.004, n = 28, Wilcoxon signed rank test, **Figure 2B**). Specifically, it increased on average by 26.9 ± 8.26 % per day across temporal EEG channels ipsilateral to the epileptogenic mesial temporal lobe (ipsi-lesional or ipsilateral). In contrast, there was no significant change of the aperiodic exponent in NEE patients (4.9 ± 2.69, p = 0.10, n = 25, Wilcoxon signed rank test; averaged across bitemporal channels). The difference between TLE and NEE patients was also significant (p = 0.019, Wilcoxon rank sum test). The effect remained significant when also using the average of unilateral channels only in NEE patients (p = 0.035, Wilcoxon rank sum test).

**Figure 2.**
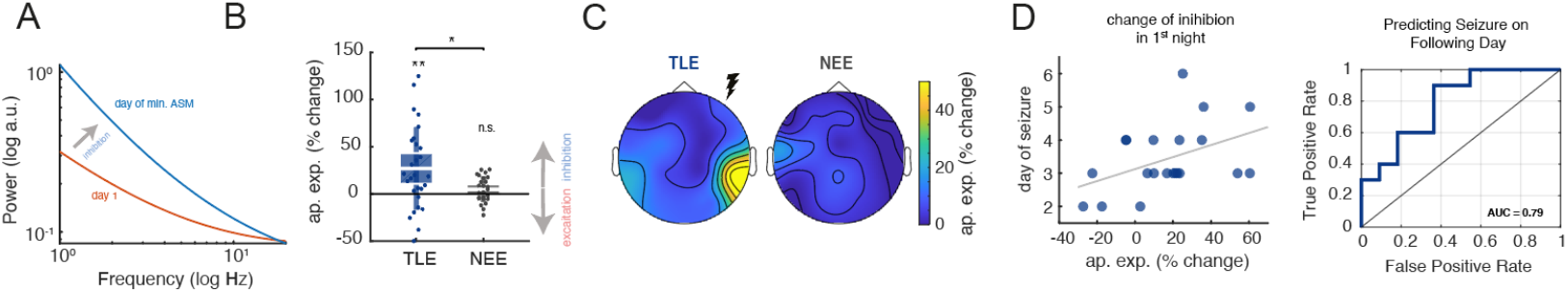
Increase in the aperiodic exponent during ASM tapering and its predictive value for seizure occurrence. **(A)** Comparison of fitted 1/fx functions for a representative patient during the initial (red) and minimum-ASM recording (blue). The steeper slope during the minimum-ASM recording indicates an increase in the aperiodic exponent, suggestive of heightened inhibitory tone. **(B)** Distributions across patients showing the average percent change in the aperiodic exponent per day for TLE (blue, left) and NEE (gray, right) patients. Note the significant change in the TLE cohort only (p = 0.004) and the significant difference between TLE and NEE patients (p = 0.019). **(C)** Topographic maps of the average aperiodic exponent change per day across patients for TLE (left) and NEE patients (right). Note the localized increase in the *ipsilesional* temporal lobe of unilateral mesial TLE patients. **(D)** Predictive analysis of seizure occurrence. Left: Correlation between the initial overnight increase in the aperiodic exponent and latency to seizures (r = 0.45, p = 0.041). Right: Receiver operating characteristic (ROC) curve for predicting seizures on the following day, with an AUC of 0.79.

The increase of the aperiodic exponent was further highly localized to the ipsilesional temporal lobe in TLE patients while no topographic hotspot could be detected in NEE patients (**Figure 2C**).

Taken together, ASM tapering is associated with a significant and highly localized increase of the aperiodic exponent in the EEG of TLE patients, indicative of a local increase of inhibitory tone.

We next explored, if this change is informative about clinical seizure activity in a predictive manner. We therefore correlated the initial increase of the aperiodic exponent over the patients first night (i.e. following the patients’ first resting state measurement to the next day) with the latency to seizures. Patients with an early seizure during this period were not considered. We indeed found a significant and positive correlation (r = 0.45, p = 0.041, **Figure 2D**), suggesting a link between a stronger increase in inhibition with later seizure occurrence. To assess the predictive value of this change, we conducted a receiver operating characteristic (ROC) analysis. Our goal was to determine how well a specific change in the aperiodic exponent during the patient’s first night (i.e., following their first RM) could predict seizure onset. The analysis revealed a promising predictive accuracy, with an area under the curve (AUC) of 0.79. The analysis identified an optimal threshold of a 7.4% increase in the aperiodic exponent for high sensitivity, achieving a sensitivity of 0.90 and a specificity of 0.64.

Taken together, the increase of the aperiodic exponent over a single night during ASM-tapering correlates with seizure-latency and can predict the occurrence of a seizure on the next day with a reasonable accuracy.

### Hippocampal IEDs trigger inhibitory responses in the lateral temporal cortex

We next wondered about the underlying mechanisms leading to the localized increase in inhibition in the EEG. To this end, we analyzed resting state recordings of 10 patients with mesial temporal lobe epilepsy undergoing intracranial recordings as part of their presurgical diagnostics. We focused our analysis on the hippocampus, the seizure onset zone in these patients, as well as the lateral temporal cortex, corresponding to the localized area of inhibition increase seen in the noninvasive EEG (**Figure 3A**).

**Figure 3.**
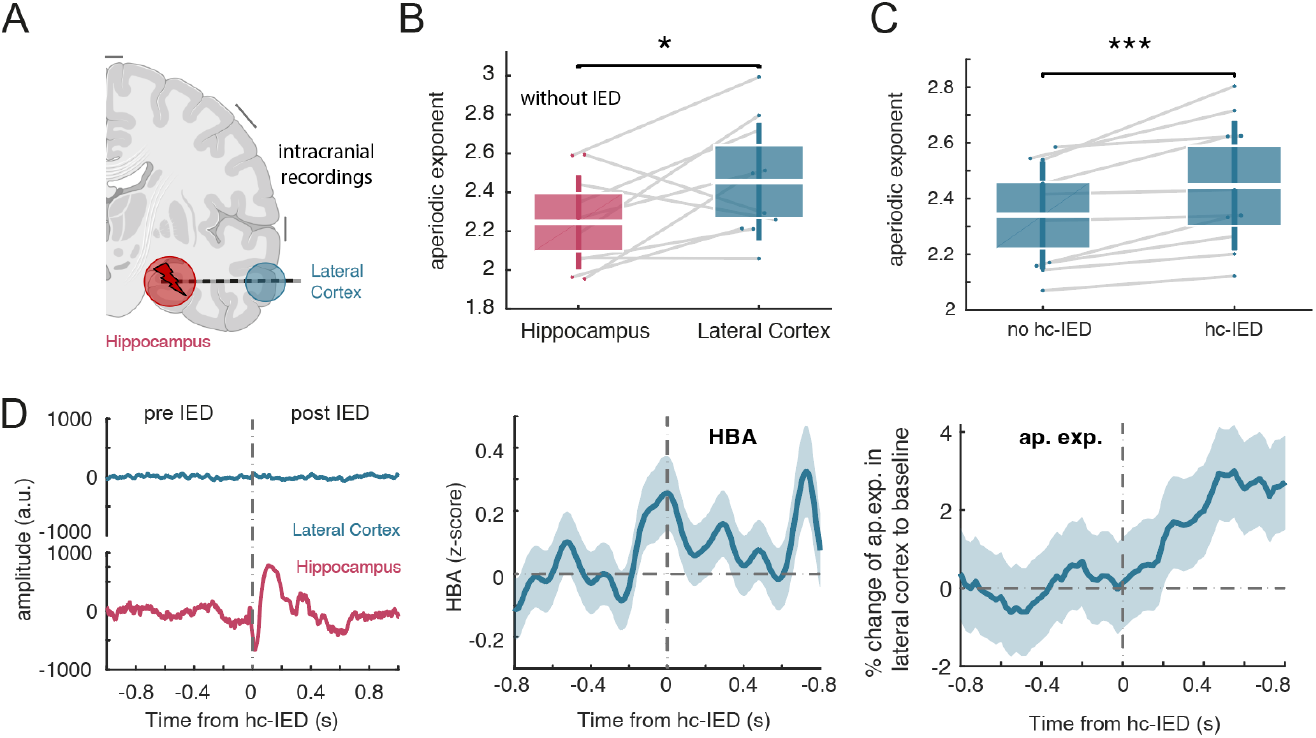
Hippocampal IEDs trigger inhibitory responses in the lateral temporal cortex. **(A)** Schematic illustration of intracranial recordings targeting the hippocampus (HC), as the seizure onset zone, and the lateral temporal cortex, corresponding to the area of increased inhibition observed in noninvasive EEG recordings. **(B)** Distributions of aperiodic exponent in the hippocampus (left, red) and lateral temporal cortex (right, blue) across patients during segments without IED (p = 0.042). **(C)** Aperiodic exponent in the lateral cortex during segments with (right) and without (left) hippocampal interictal epileptiform discharges (hc-IEDs). Note the significant increase during hc-IED segments (p = 9.8 × 10^−4^) **(D)** Time-resolved analysis of lateral cortical responses triggered by hc-IEDs. Left: Representative single trial traces of hippocampal and lateral cortical activity pre- and post-hc-IED. Middle: High-frequency band activity (HBA, z-score) in the lateral cortex shows a significant transient increase following hc-IEDs (p = 2.19 × 10^−55^). Right: Aperiodic exponent changes in the lateral cortex post-hc-IEDs reveal a significant transient increase in synaptic inhibition (p = 0.006). Shaded areas depict SEM.

First, we compared the spontaneous aperiodic exponent for both areas (**Figure 3B**). We found the average aperiodic exponent to be significantly higher in lateral cortex across patients (p = 0.042, n = 10, Wilcoxon signed rank test). Specifically, the aperiodic exponent was on average 2.14 ± 0.06 in the hippocampus and 2.34 ± 0.06 in the lateral cortex. Thus, the aperiodic exponent was about 9% higher in the lateral cortex, indicating higher synaptic inhibition compared to the hippocampus.

But what might be the cause of this higher inhibitory tone in the lateral temporal cortex? Given the tight connectivity between the hippocampus and the lateral temporal cortex, it appears conceivable that the increase in inhibitory tone might be secondary to the ongoing epileptic activity and associated increased excitability in the hippocampus. We therefore wondered, if interictal epileptic activity in the form of hippocampal interictal epileptiform discharges (hc-IED) might be related to a change of the aperiodic exponent in the lateral cortex. To this end, we compared the aperiodic exponent in the lateral cortex in 3s segments during which hc-IED occurred with those where no hc-IED were present (**Figure 3C**). Indeed, the aperiodic exponent was significantly higher during segments with hc-IED (p = 9.8 × 10^−4^, n = 10, averaged across patients). Specifically, the aperiodic exponent in the lateral cortex was 2.45 ± 0.07 in segments with hc-IED and 2.34 ± 0.06 in segments without hc-IED.

To relate changes in the lateral cortex more directly to hc-ED, we triggered the lateral cortex’ signal on hc-IED occurrence (**Figure 3D**). First, we analyzed the hc-IED’s effect on neuronal activity in the lateral cortex. Given that hc-IEDs represent abrupt, highly synchronized neuronal firing in the hippocampus, we hypothesized that this activity might influence neuronal firing in the lateral cortex, even in the absence of local IED there. To test this, we utilized high-frequency band activity (HBA) as a proxy for neuronal activity (Helfrich et al. 2018), as standard clinical intracranial macroelectrodes cannot capture single- or multi-unit activity directly. Consistent with our hypothesis, we observed a significant transient increase in HBA, accompanied by a sustained component, following hc-IEDs (p = 2.19 × 10^−55^, [0-0.8s] Wilcoxon signed rank test; **Figure 3D, middle panel**). Thus hc-IED trigger transient firing in the lateral cortex.

Next, we investigated whether hc-IEDs also induce changes in the aperiodic exponent in the lateral cortex, as a marker of altered synaptic inhibition in connected regions. Indeed, we found a significant increase in the aperiodic exponent in the lateral cortex following hc-IEDs (p = 0.006, Wilcoxon signed-rank test**; Figure 3D, right panel**).

Thus, hc-IED lead to a transient increase in neuronal firing in the lateral cortex associated with an increase in synaptic inhibition. This mechanism may also contribute to the elevated inhibitory tone observed in the lateral cortex and the EEG: repetitive IED induce transient inhibitory responses that, over time, could result in a sustained increase in inhibitory tone (as reflected in periods without IED, **Figure 3B**, and EEG **Figure 2C**) through plasticity-related processes.

### Hippocampal IEDs alter directed connectivity between the Hippocampus and Lateral Cortex

In a last step, we investigated how hc-IEDs influence the directed flow of information between the hippocampus and the lateral cortex (**Figure 4A**). This analysis was grounded in the hypothesis that the seizure onset zone, as the origin of propagating epileptic activity, would exert a stronger influence on surrounding areas than it would receive in return. To test this hypothesis, we applied Granger causality analysis to examine the directional interactions between the hippocampus and the lateral cortex. In brief, Granger causality quantifies the extent to which activity in one region helps predict the future activity in another, thereby providing a robust measure of the directional influence and information flow between interconnected brain regions.

**Figure 4.**
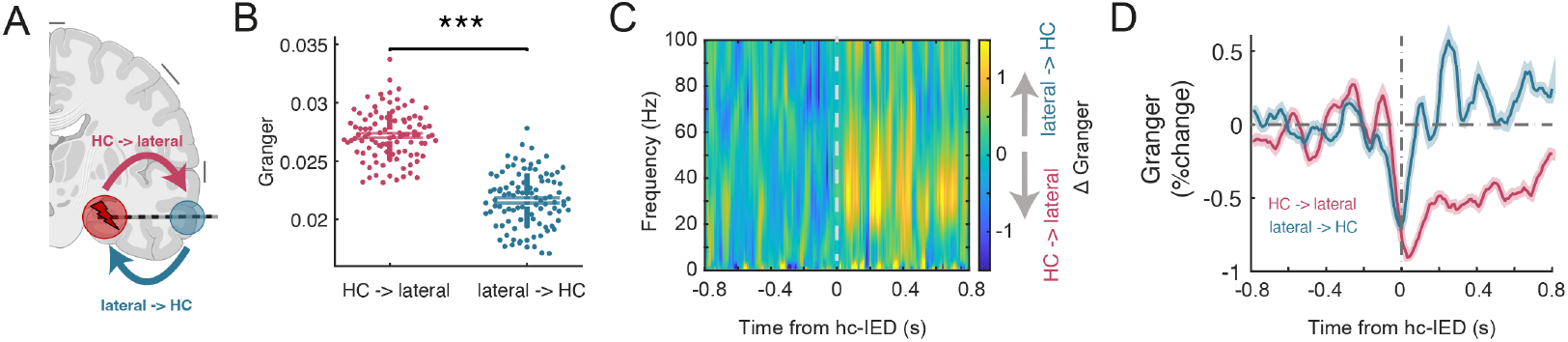
IED disrupt hippocampal dominance over the lateral cortex. **(A)** Schematic representation of directional connectivity between the hippocampus (HC) and lateral cortex, highlighting HC→lateral (red) and lateral→HC (blue) pathways. **(B)** Baseline Granger causality distributions (boxplot) reveal significantly stronger HC→lateral connectivity (right, left) compared to lateral→HC connectivity (blue, right) prior to hc-IEDs (p = 6.4×10^−30^, Wilcoxon rank sum test). Dots represent surrogate trials. **(C)** Differential time-frequency analysis (peri-hc-IED Granger causality: lateral→HC - HC→lateral) shows changes in Granger causality induced by hc-IEDs. Note the increase for the lateral→HC direction in the low gamma range (20-50 Hz). **(D)** Time-resolved analysis of Granger causality changes averaged in the low gamma band (20-50 Hz) following hc-IEDs for the HC→lateral direction (red) and lateral→HC direction (blue). Shaded areas depict SEM based on surrogate statistics.

First, we compared Granger causality during baseline conditions (i.e., prior to hc-IEDs). Consistent with our hypothesis, we observed significantly higher Granger causality values for hippocampus-to-lateral cortex connectivity (HC→lateral) compared to the reverse direction (lateral→HC) (p = 6.4×10^−30^, Wilcoxon rank sum test and surrogate statistics). This supports the notion that the hippocampus, as the seizure onset zone, exerts a stronger influence on the lateral cortex under baseline conditions.

We next explored how hc-IEDs influence directed connectivity between the hippocampus and the lateral cortex. A differential time-frequency analysis (**Figure 4C**) highlighted a pronounced change in the low gamma range (20–50 Hz). We therefore focused on this frequency band to yield a more detailed, time-resolved representation (**Figure 4D**). The analysis revealed a significant reduction in HC→lateral connectivity following hc-IED (mean change: -0.52%; 95% CI: [-0.594, -0.446], p < 0.01, surrogate statistics). In contrast, lateral→HC connectivity exhibited a significant increase (mean change: 0.15%; 95% CI: [0.006, 0.316], p = 0.04, surrogate statistics). The observed difference in connectivity changes between these two directions was statistically significant (p < 0.01). Importantly, these changes reflect average effects induced by single hc-IEDs.

These findings suggest that hc-IEDs disrupt the hippocampus’s dominant influence over the lateral cortex, altering the balance of connectivity dynamics.

## Discussion

This study investigates the dynamics of E/I balance in mesial temporal lobe epilepsy, leveraging a recently developed and validated method to quantify inhibitory dynamics and their regional variability. Contrary to the view that epilepsy is characterized predominantly by excessive neuronal excitability, our findings reveal that TLE involves complex, region-specific inhibitory mechanisms, particularly during interictal periods and antiseizure medication tapering.

### Localized inhibition and predictive value of the aperiodic exponent

We found a significant and localized increase in the aperiodic exponent in the ipsilesional temporal lobe of TLE patients during ASM tapering using EEG, indicative of heightened inhibitory tone. This effect was absent in NEE controls. These findings are in line with recent results reporting higher aperiodic exponents in epilepsy patients compared to healthy controls (Duma et al. 2024). In our case, the increase in inhibition was highly localized to the ipsilesional temporal cortex, potentially identifying an “inhibitory zone” closely related to the epileptogenic zone. How this potential “inhibitory zone” relates to other zones (Rosenow and Lüders 2001), particular the irritative zone (i.e. areas with IED), will need to be addressed in future studies.

Using intracranial recordings, we further observed that the lateral temporal cortex exhibited significantly higher aperiodic exponents compared to the hippocampus, suggesting stronger synaptic inhibition in the lateral cortex. This regional difference highlights a functional differentiation: while the seizure onset zone might show increased excitability, eventually triggering seizures, surrounding regions such as the lateral cortex exhibit increased inhibitory tone, possibly as an adaptive mechanism to contain epileptic activity. This observation complements our EEG findings and extends the concept of an “inhibitory penumbra”, previously described in experimental and in vitro studies during seizure propagation (e.g. Trevelyan et al. 2006; Prince and Wilder 1967; Schevon et al. 2012), to the interictal period.

Further, we found that the increase in inhibition following ASM reduction correlated with seizure latency during VEM and predicted seizure occurrence on the subsequent day. This provides a potential biomarker for predicting seizure occurrence during presurgical monitoring, emphasizing the clinical relevance of interictal inhibitory dynamics.

### The role of hippocampal IEDs in neocortical inhibition

How does this increase in inhibition come about? Our results reveal that hc-IED trigger transient increases in inhibition in the lateral temporal cortex. These inhibitory responses were accompanied by heightened neuronal firing, reflected in high-frequency band activity. Such transient but repetitive responses may contribute to the sustained elevation of inhibitory tone observed in the lateral cortex and the EEG during ASM tapering, potentially via plasticity-related processes. Yet, whether the hc-IED-triggered increase in inhibition is mediated through direct activation of interneurons or via secondary network effects needs further investigation. One possibility is that hc-IEDs directly activate interneurons in the lateral temporal cortex through monosynaptic pathways. This hypothesis aligns with evidence showing that hippocampal outputs can strongly influence downstream cortical interneurons (e.g. (Tierney et al. 2004; Thierry et al. 2000)). Alternatively, the observed inhibition could arise via secondary network effects, where excitatory activity in the hippocampus indirectly drives inhibitory circuits in the lateral cortex through polysynaptic pathways. Plasticity-related mechanisms, such as long-term potentiation of inhibitory synapses, may further amplify these responses over time.

### Effect of hc-IED on directional connectivity

Finally, Granger causality analysis demonstrated that hc-IED disrupt the hippocampus’s dominant influence over the lateral cortex. Specifically, we observed a reduction in hippocampus-to-lateral cortex connectivity and a concurrent increase in lateral cortex-to-hippocampus connectivity following hc-IEDs. This reversal in connectivity dynamics suggests a disruption of hierarchical control, potentially reflecting compensatory network responses to hippocampal hyperexcitability. Changed connectivity patterns are a recurring feature in epilepsy, as previous studies have reported directional shifts during seizures within the hippocampus (Cadotte et al., 2010) and altered global connectivity in epilepsy patients compared to healthy controls (Ji et al., 2013). Our findings extend these phenomena to the interictal period and hippocampal-cortical interactions with a dynamical reversal of the connectivity pattern.

### Limitations and future directions

While our study provides valuable insights, several limitations should be acknowledged. First, the use of ASM tapering as a modulator of E/I balance introduces confounding factors related to withdrawal-induced changes in network activity. However, since both the TLE and control groups underwent ASM tapering, it is unlikely that nonspecific withdrawal effects alone account for the TLE-specific changes we observed. Second, while the aperiodic exponent appears to serve as a valid marker of synaptic inhibition (Diehl and Redish 2024; Pochinok et al. 2024; Brake et al. 2024), its exact physiological correlates remain to be fully understood (see also (Schmidt et al. 2024). Complementary studies using optogenetic or pharmacological approaches could further validate the link between aperiodic exponent changes and inhibitory tone.

Additionally, our findings are limited to mesial TLE and may not generalize to other epilepsy subtypes. Future studies will need to test whether similar inhibitory mechanisms are present in other focal or even generalized epilepsies. Finally, while we demonstrated the predictive value of the aperiodic exponent for seizure occurrence, larger cohorts and prospective studies are necessary to validate its clinical utility as a biomarker.

### Conclusions

In summary, this study provides evidence that TLE is characterized by complex, region-specific inhibitory mechanisms extending beyond the seizure onset zone. By integrating scalp and intracranial EEG data, we demonstrate that interictal inhibitory responses in the lateral temporal cortex are driven by hippocampal IEDs and can be quantified non-invasively using the aperiodic exponent. These findings refine our understanding of the E/I balance in epilepsy and elucidate potential mechanisms that could guide future approaches to study epileptic networks, responses to ASM and managing seizure activity.

## Methods

### Recordings and data pre-processing

We included EEG recordings from 28 patients with unilateral mesial temporal lobe epilepsy and 25 patients with non-epileptic episodes only (NEE) who underwent a video-EEG monitoring (VEM) with tapering of anti-seizure medication (ASM) at a tertiary epilepsy center in Frankfurt, Germany. Additionally, 10 TLE patients undergoing invasive video-EEG monitoring with mesio-temporal coverage and a hippocampal irritative zone and seizure onset zone were included to study intracranial dynamics. Patients in both the TLE and NEE cohorts received similar treatment and underwent similar diagnostic procedures. Screened patients with recordings affected by excessive artifacts were excluded from the analysis. Given the rare characteristics of this cohort, the patients’ data has – in part – already been included in previous studies for other scientific questions (Kienitz et al. 2024).

The study received approval from the Ethics Committee of the Medical Faculty at Goethe University Frankfurt. Informed consent for data analysis was waived due to the retrospective design of the study. Clinical and demographic information was obtained from digital patient records.

EEG recording and preprocessing has previously been described (Kienitz et al. 2024). In short, EEG was recorded as resting state once daily according to the 10-20 system with few extensions according to the 10-10 system (e.g. FT9/FT10). EEG was recorded using the BRAIN QUICK soft- and hardware (Micromed, Mogliano Veneto, Italy). EEG electrodes were gold-plated (Au)-electrodes. Fpz and an additional electrode positioned along the midline between Fz and Cz were used as reference electrodes. EEG data was visually inspected, and segments containing open eyes, interictal epileptiform discharges (IED), or artifacts (biological or technical) were excluded. The data was bandpass-filtered (0.5–40 Hz, 2nd-order Butterworth filter) and segmented into 3-second epochs for analysis. Notch-filters were used to reduce line noise.

Intracranial recordings were conducted using sEEG Depth Electrodes (Ad-Tech Medical) on clinical indication. Data was low-pass filtered at 500 Hz and down-sampled to 1 kHz

We focused our analysis on one electrode contact within in the hippocampus and in the lateral temporal cortex of the same electrode, as validated by MRI location and typical electrophysiological signals. Both were re-referenced to a contact in the white matter. IED were detected as described previously (Kienitz et al. 2024). In short, the data was transformed into velocity space, where outliers were identified as deviations exceeding three standard deviations from a moving median calculated over a 15-second window. Following the method described by Gelinas et al., an additional amplitude criterion was then applied, requiring the IED amplitude to exceed the pre-IED range by a factor of two (Gelinas et al. 2016). IED occurring within 1 second before or after another IED were excluded.

The analyses were performed using custom MATLAB scripts (MathWorks, Inc., Natick, MA, USA) and the FieldTrip toolbox (Oostenveld et al. 2011).

### Computation of the aperiodic exponent

To quantify inhibitory dynamics, the aperiodic exponent was computed for each EEG 3s segment using the multitaper method in the frequency range of 1-20 Hz and the FOOOF algorithm implemented in FieldTrip. This way, power spectra were decomposed into periodic and aperiodic components, and the slope of the 1/f component was extracted as the aperiodic exponent. We calculated the aperiodic exponent across all available EEG channels for each patient. The average changes per day were assessed using a linear regression model between the first resting state measurement and the minimum ASM day. Topographic maps of the aperiodic exponent were generated using FieldTrip to visualize spatial patterns of inhibitory dynamics. Data from TLE and NEE cohorts were contrasted to identify group differences.

To assess the predictive accuracy of the aperiodic exponent, we trained general linear regression models and evaluated their performance using ROC curves. The area under the curve (AUC) provided a measure of classification accuracy between TLE and NEE patients. For the ROC and the correlational analysis with seizure latency, the change of the aperiodic exponent between the first resting state measurement and the consecutive day was used.

### Analysis of intracranial data

For the analysis underlying **Figure 3B-C** we cut the intracranial data from the hippocampal and lateral cortical contact into 3s segments and categorized them according to the presence of IED on the hippocampal contact (hc-IED). In **Figure 3B** we compared the average aperiodic exponent across segments *without* hc-IED between the hippocampal and the lateral cortical contact. For **Figure 3C**, we compared segments of the lateral cortical contacts *with and without* concurrent hc-IED.

In a next step, we triggered the lateral cortical signal on hc-IED occurrence, spanning a peri-event window of -1 to 1 seconds, and pooled these trials across patients. Given potential filtering artifacts at the borders of the analysis window, we eventually focused on the time range of -0.8 to 0.8 s.

To assess hc-IED effects on neuronal activity in the lateral cortex, we made use of high-frequency band activity (HBA) which serves as a proxy for local neuronal firing. HBA was computed as described in (Helfrich et al. 2018). For each trial, a bandpass filter was applied to the data in overlapping frequency bins of 10 Hz within the HBA range of 70–150 Hz. The Hilbert transform was then applied to compute the analytic amplitude (envelope) of the filtered signal. The envelopes were averaged across frequency bins to provide a single HBA time series per trial. A baseline period from -0.8 to -0.2 s relative to IED onset was used to normalize the HBA using a z-scoring. Before averaging across trials, outliers were automatically removed.

To estimate the aperiodic exponent in a peri-hc-IED time-resolved manner, we computed the aperiodic exponent of the lateral cortical data in sliding time windows of 50 ms length which was shifted in steps of 25 ms. The computation of the aperiodic exponent of intracranial data focused on the frequency range of 30-100 Hz. Baseline normalization was performed as percent change relative to a baseline period of -0.8 to -0.2 s. Before averaging across trials, outliers were automatically removed.

### Granger Causality Analysis

We performed Granger causality analysis (Granger 1969) to examine directional interactions between hippocampal and lateral temporal regions in the context of hc-IED using the Fieldtrip implementation of Granger causality. First, multivariate autoregressive (MVAR) modeling was applied using a sliding time window of 50 ms, with a time resolution of 10 ms and frequency resolution spanning from 1 to 100 Hz. Directional Granger causality estimates were computed for mesio-to-lateral and lateral-to-mesio interactions. **Figure 4C** displays the difference of Granger causality values of both directions: “lateral→HC” – “HC→lateral”. For **Figure 4D**, Granger causality values were averaged across frequencies in the frequency range between 20 and 50 Hz.

To account for variability across trials, a bootstrap resampling approach was employed. For each of 100 bootstrap iterations, trials were sampled with replacement to construct surrogate datasets of equal size. Granger causality was computed for each bootstrap iteration, yielding surrogate distributions of connectivity estimates. Bootstrap distributions were used to compute 95% confidence intervals (CI) for each condition and for the difference between conditions (mesio-to-lateral vs. lateral-to-mesio). P-values were derived using a two-tailed test. Connectivity values were normalized relative to a baseline period (-0.8 to -0.2 seconds) as percent change.

## Acknowledgments

Figures 1A, 3A and 4A were created with BioRender.com.

## References

Abdallah, Chifaou, Daniel Mansilla, Erica Minato, Christophe Grova, Sandor Beniczky, and Birgit Frauscher. 2024. “Systematic Review of Seizure-Onset Patterns in Stereo-Electroencephalography: Current State and Future Directions.” Clinical Neurophysiology 163 (July):112–23. 10.1016/j.clinph.2024.04.016.

Ahmad, Jumana, Claire Ellis, Robert Leech, Bradley Voytek, Pilar Garces, Emily Jones, Jan Buitelaar, et al. 2022. “From Mechanisms to Markers: Novel Noninvasive EEG Proxy Markers of the Neural Excitation and Inhibition System in Humans.” Translational Psychiatry 12 (1):467. 10.1038/s41398-022-02218-z.

Brake, Niklas, Flavie Duc, Alexander Rokos, Francis Arseneau, Shiva Shahiri, Anmar Khadra, and Gilles Plourde. 2024. “A Neurophysiological Basis for Aperiodic EEG and the Background Spectral Trend.” Nature Communications 15 (1):1514. 10.1038/s41467-024-45922-8.

Diehl, Geoffrey W., and A. David Redish. 2024. “Measuring Excitation-Inhibition Balance through Spectral Components of Local Field Potentials.” bioRxiv. 10.1101/2024.01.24.577086.

Donoghue, Thomas, Matar Haller, Erik J. Peterson, Paroma Varma, Priyadarshini Sebastian, Richard Gao, Torben Noto, et al. 2020. “Parameterizing Neural Power Spectra into Periodic and Aperiodic Components.” Nature Neuroscience 23 (12):1655–65. 10.1038/s41593-020-00744-x.

Duma, Gian Marco, Simone Cuozzo, Luc Wilson, Alberto Danieli, Paolo Bonanni, and Giovanni Pellegrino. 2024. “Excitation/Inhibition Balance Relates to Cognitive Function and Gene Expression in Temporal Lobe Epilepsy: A High Density EEG Assessment with Aperiodic Exponent.” Brain Communications 6 (4):fcae231. 10.1093/braincomms/fcae231.

Fisher, R. S., W. R. Webber, R. P. Lesser, S. Arroyo, and S. Uematsu. 1992. “High-Frequency EEG Activity at the Start of Seizures.” Journal of Clinical Neurophysiology: Official Publication of the American Electroencephalographic Society 9 (3):441–48. 10.1097/00004691-199207010-00012.

Gao, Richard, Erik J. Peterson, and Bradley Voytek. 2017. “Inferring Synaptic Excitation/Inhibition Balance from Field Potentials.” NeuroImage 158 (September):70–78. 10.1016/j.neuroimage.2017.06.078.

Gelinas, Jennifer N, Dion Khodagholy, Thomas Thesen, Orrin Devinsky, and György Buzsáki. 2016. “Interictal Epileptiform Discharges Induce Hippocampal-Cortical Coupling in Temporal Lobe Epilepsy.” Nature Medicine 22 (6):641–48. 10.1038/nm.4084.

Granger, C. W. J. 1969. “Investigating Causal Relations by Econometric Models and Cross-Spectral Methods.” Econometrica 37 (3):424–38. 10.2307/1912791.

He, Biyu J. 2014. “Scale-Free Brain Activity: Past, Present, and Future.” Trends in Cognitive Sciences 18 (9):480–87. 10.1016/j.tics.2014.04.003.

Helfrich, Randolph F., Ian C. Fiebelkorn, Sara M. Szczepanski, Jack J. Lin, Josef Parvizi, Robert T. Knight, and Sabine Kastner. 2018. “Neural Mechanisms of Sustained Attention Are Rhythmic.” Neuron 99 (4):854–65. 10.1016/j.neuron.2018.07.032.

Huberfeld, Gilles, Lucia Wittner, Stéphane Clemenceau, Michel Baulac, Kai Kaila, Richard Miles, and Claudio Rivera. 2007. “Perturbed Chloride Homeostasis and GABAergic Signaling in Human Temporal Lobe Epilepsy.” Journal of Neuroscience 27 (37):9866–73. 10.1523/JNEUROSCI.2761-07.2007.

Kienitz, Ricardo, Michael Strüber, Nina Merkel, Annika Süß, Andrea Spyrantis, Adam Strzelczyk, and Felix Rosenow. 2024. “Neuronal Complexity Tracks Changes of Epileptic Activity and Identifies Epilepsy Patients Independent of Interictal Epileptiform Discharges.” Epilepsia, in Press.

Lanerolle, N. C. de, J. H. Kim, R. J. Robbins, and D. D. Spencer. 1989. “Hippocampal Interneuron Loss and Plasticity in Human Temporal Lobe Epilepsy.” Brain Research 495 (2):387–95. 10.1016/0006-8993(89)90234-5.

Marx, Markus, Carola A. Haas, and Ute Häussler. 2013. “Differential Vulnerability of Interneurons in the Epileptic Hippocampus.” Frontiers in Cellular Neuroscience 7. 10.3389/fncel.2013.00167.

McCormick, D. A., and D. Contreras. 2001. “On the Cellular and Network Bases of Epileptic Seizures.” Annual Review of Physiology 63:815–46. 10.1146/annurev.physiol.63.1.815.

Ngugi, Anthony K., Christian Bottomley, Immo Kleinschmidt, Josemir W. Sander, and Charles R. Newton. 2010. “Estimation of the Burden of Active and Life-Time Epilepsy: A Meta-Analytic Approach.” Epilepsia 51 (5):883–90. 10.1111/j.1528-1167.2009.02481.x.

Oostenveld, Robert, Pascal Fries, Eric Maris, and Jan-Mathijs Schoffelen. 2011. “FieldTrip: Open Source Software for Advanced Analysis of MEG, EEG, and Invasive Electrophysiological Data.” Computational Intelligence and Neuroscience, January. 10.1155/2011/156869.

Pochinok, Irina, Tristan M. Stöber, Jochen Triesch, Mattia Chini, and Ileana L. Hanganu-Opatz. 2024. “A Developmental Increase of Inhibition Promotes the Emergence of Hippocampal Ripples.” Nature Communications 15 (1):738. 10.1038/s41467-024-44983-z.

Prince, David A., and B. Joe Wilder. 1967. “Control Mechanisms in Cortical Epileptogenic Foci*: ‘Surround’ Inhibition.” Archives of Neurology 16 (2):194–202. 10.1001/archneur.1967.00470200082007.

Rosenow, Felix, and Hans Lüders. 2001. “Presurgical Evaluation of Epilepsy.” Brain 124:1683–1700. 10.4103/1817-1745.40593.

Schevon, Catherine a., Shennan a. Weiss, Guy McKhann, Robert R. Goodman, Rafael Yuste, Ronald G. Emerson, and Andrew J. Trevelyan. 2012. “Evidence of an Inhibitory Restraint of Seizure Activity in Humans.” Nature Communications 3:1060. 10.1038/ncomms2056.

Schmidt, Fabian, Sarah K. Danböck, Eugen Trinka, Dominic P. Klein, Gianpaolo Demarchi, and Nathan Weisz. 2024. “Age-Related Changes in ‘Cortical’ 1/f Dynamics Are Linked to Cardiac Activity.” bioRxiv. 10.1101/2022.11.07.515423.

Spencer, Susan S. 2002. “Neural Networks in Human Epilepsy: Evidence of and Implications for Treatment.” Epilepsia 43 (3):219–27. 10.1046/j.1528-1157.2002.26901.x.

Staley, Kevin. 2015. “Molecular Mechanisms of Epilepsy.” Nature Neuroscience 18 (3):367–72. 10.1038/nn.3947.

Thierry, Anne-Marie, Yves Gioanni, Eric Dégénétais, and Jacques Glowinski. 2000. “Hippocampo-Prefrontal Cortex Pathway: Anatomical and Electrophysiological Characteristics.” Hippocampus 10 (4):411–19. 10.1002/1098-1063(2000)10:4<411::AID-HIPO7>3.0.CO;2-A.

Tierney, Patrick L., Eric Dégenètais, Anne-Marie Thierry, Jacques Glowinski, and Yves Gioanni. 2004. “Influence of the Hippocampus on Interneurons of the Rat Prefrontal Cortex.” European Journal of Neuroscience 20 (2):514–24. 10.1111/j.1460-9568.2004.03501.x.

Trevelyan, Andrew J., David Sussillo, Brendon O. Watson, and Rafael Yuste. 2006. “Modular Propagation of Epileptiform Activity: Evidence for an Inhibitory Veto in Neocortex.” Journal of Neuroscience 26 (48):12447–55. 10.1523/JNEUROSCI.2787-06.2006.

